# PLAPT: Protein-Ligand Binding Affinity Prediction Using Pretrained Transformers

**DOI:** 10.1101/2024.02.08.575577

**Authors:** Tyler Rose, Nicolò Monti, Navvye Anand, Tianyu Shen

**Affiliations:** Wolfram Research, Cupertino, CA; ASC27, Pescara, IT; Sanskriti School, New Delhi, IN; Newport High School, Bellevue, WA

**Keywords:** Protein-Ligand Binding Affinity, Transformer Models, Predictive Biology, Drug Discovery

## Abstract

Predicting protein-ligand binding affinity is crucial for drug discovery, as it enables efficient identification of drug candidates. We introduce PLAPT, a novel model utilizing transfer learning from pre-trained transformers like ProtBERT and ChemBERTa to predict binding affinities with high accuracy. Our method processes one-dimensional protein and ligand sequences, leveraging a branching neural network architecture for feature integration and affinity estimation. We demonstrate PLAPT’s superior performance through validation on multiple datasets, achieving state-of-the-art results while requiring significantly less computational resources for training compared to existing models. Our findings indicate that PLAPT offers a highly effective and accessible approach for accelerating drug discovery efforts.

## 1 Introduction

The process of drug discovery hinges significantly on understanding the interactions between proteins and ligands, particularly the measure of protein-ligand binding affinity. This metric, indicative of the interaction strength between a protein (the target) and a ligand (a smaller molecule capable of binding to the target), is foundational in identifying promising drug candidates. Binding affinities are quantitatively expressed as dissociation constants (*K*_*d*_), where lower values denote stronger and more favorable bindings. Accurate binding affinities prediction through *in silico* methods reduce web-lab testing, which is significant in accelerating drug development [1]. This efficiency is particularly vital during global pandemics, allowing for rapid identification of promising drug candidates, conserving time and resources, and enabling a swift response to disease outbreaks.

Traditionally, the prediction of protein-ligand binding affinities heavily relied on physics-based computational methods with molecular dynamics programs being at the forefront [2, 3, 4]. However, these methods can face accuracy issues due to simplifications and poor quality-scoring functions, leading to discrepancies between predicted and experimental binding affinities [5]. Despite these limitations, the foundational role of physics-based methods in understanding protein-ligand interactions cannot be understated. They paved the way for the integration of computational tools in drug discovery, setting the stage for the emergence of machine learning and deep learning approaches. These newer methodologies promised to overcome scalability and accuracy challenges by leveraging advanced learning algorithms to predict binding affinities [6].

The transition towards machine learning, and subsequently deep learning, marked a significant evolution in the computational prediction of binding affinities. Deep learning, in particular, have shown great promise in autonomously learning complex features from raw data [7]. These methods are broadly categorized into structure-based and sequence-based approaches. Structure-based methods, such as Pafnucy and OnionNet, utilize three-dimensional (3D) structural information of proteins and ligands to predict binding affinities [8, 9]. Conversely, sequence-based methods, such as DeepDTAF, CAPLA, and DTITR rely on one-dimensional (1D) sequences, which are simpler and have exhibited similar performances to their 3D counterparts [10, 11, 12]. The emergence of transformer models has demonstrated the remarkable capacity of machine learning models to analyze and comprehend one-dimensional sequential data [13]. This advancement has positioned transformers as an integral component in the contemporary landscape of bioinformatics, specifically within the domains of protein-sequence and molecular-sequence modeling [14, 15, 16].

Transformer models have naturally emerged as a groundbreaking approach in leveraging sequence data to predict protein-ligand binding affinity. While models like CAPLA utilize a cross-attention mechanism to achieve high accuracy, they do not leverage the transfer learning benefits provided by pre-trained encoders [17, 11]. By employing pre-trained encoder models trained unsupervised on large datasets, Blanchard et al. produced a state-of-the-art model with impressive generalization capabilities [18]. However, their approach is extremely computationally intensive [18], and we found that comparable or higher accuracy can be achieved using a much simpler prediction model with computational resources five orders of magnitude less to that of the existing approach.

Against this backdrop, our research introduces the Protein Ligand Binding Affinity Prediction Using Pre-trained Transformers (PLAPT) model. PLAPT represents a novel contribution to the field by efficiently integrating knowledge encapsulated in pre-trained models such as ProtBERT and ChemBERTa [19, 16], while using minimal computational resources to achieve state-of-the-art accuracy. Through a branching neural network (BNN) architecture, PLAPT integrates features from both protein and ligand embeddings to estimate binding affinity values with high precision. This is evident by our performance metric analysis, comparing PLAPT’s accuracy with that of other protein-ligand prediction models. Additionally, similar to the Blanchard et al. model, PLAPT does not include explicit protein pocket information [18]. This decision is based on the substantial representational capacity of the ProtBERT encoder and the simplified data acquisition process made possibly by only considering entire amino-acid sequences.

PLAPT’s innovation lies in its ability to achieve high performance levels while being trained with significantly lower computational and data requirements compared to previous models. This efficiency is primarily due to the use of open-source pre-trained encoders, significantly reducing the model’s dependence on extensive computational resources. By demonstrating that using less compute and data for training can still yield state-of-the-art results, PLAPT’s methods pave the way for further innovations and rapid iteration in drug discovery research, improving accessibility and efficiency.

## 2 Methods

### 2.1 Input Representation

PLAPT is designed to work solely with one-dimensional string inputs, necessitating just a protein sequence and the SMILES notation of a ligand for making predictions. This design contrasts with models such as CAPLA, which demand data on the protein pocket [11]. Refer to Figure 1 for an illustration of the inputs used by PLAPT.

**Figure 1:**
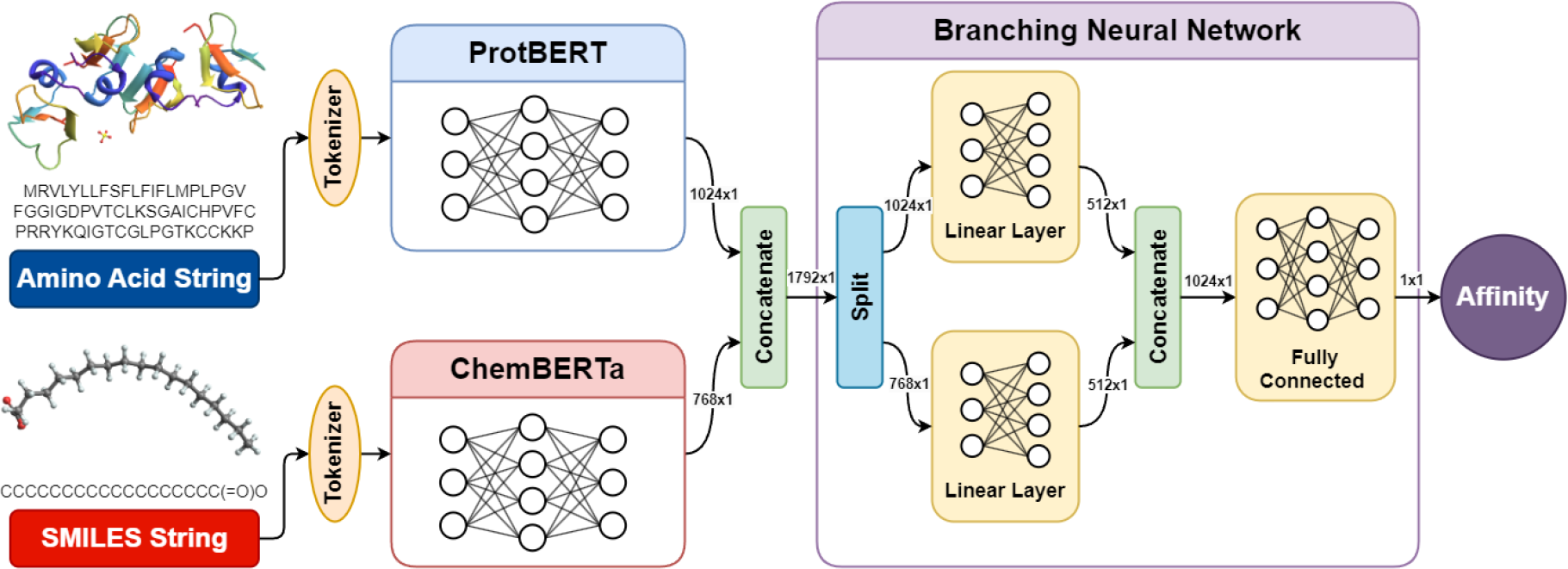
Architectural diagram of PLAPT. The protein and ligand sequence encoders are connected to a branching neural network which contains bifurcated linear layers, who’s outputs are concatenated into fully connected layers that predict binding affinity.

#### Protein Input

For the purpose of PLAPT, proteins are inputted as a single string of amino acid residues. This sequence is subject to pre-processing steps which include the separation of each character by spaces and the substitution of non-standard amino acids (e.g., B, U, Z, O, J) with the character ‘X’. Subsequently, characters are converted to tokens using the tokenization strategy described by Elnaggar et al. for ProtBERT [19, 20]. The tokenization process transforms the amino acid strings into a combination of a padded token list (length 3200) and an accompanying attention mask (length 3200), both of which are then inputted into the ProtBERT encoder [20]. It is pertinent to note that ProtBERT [19, 20] is structured to accommodate sequences with lengths up to 40,000. However, due to computational constraints inherent to transformer models when dealing with longer sequences, the length of each protein token sequence is adjusted to 3200 tokens through either padding or truncation. This modification is anticipated to have a minimal impact on the model’s predictive accuracy, as only approximately 0.2% of sequences within our dataset exceed this length threshold.

#### Molecule Input

In the case of chemical compounds, the input to PLAPT is provided in the form of canonical SMILES strings. The decision to employ SMILES representations leverages the natural language processing capabilities of transformer models, thanks to their compact and human-readable nature which facilitates easy integration with existing bioinformatics tools. Moreover, despite the absence of three-dimensional spatial data characteristic of graph-based molecule representations, the proficiency of transformer models in processing one-dimensional sequential data and recognizing intricate patterns as well as long-range dependencies within these sequences renders them apt for this application [13]. The SMILES strings are tokenized using the SmilesTokenizer developed by Schwaller et al. [21], which performs zero padding to standardize the length of all token sequences to 278. This choice of padding length corresponds to the longest token sequence encountered in our dataset, although our molecule encoder, ChemBERTa, is capable of supporting sequences up to a length of 512. Similar to the protein tokenizer, the SMILES tokenizer generates an attention mask alongside the tokenized output, for use in the ChemBERTa encoder.

### 2.2 Model Architecture

The PLAPT architecture of the PLAPT contains two primary components: a feature extraction module leveraging the capabilities of pre-trained BERT-like encoders, as well as a prediction module constituted by a novel branching neural network (BNN) design.

#### 2.2.1 Feature Extraction

Studies like CAPLA demonstrate the efficacy of a combined approach to feature extraction, including the use of a cross-attention mechanism, in enhancing predictive performance for protein-ligand interaction tasks [11]. However, leveraging the advanced biochemical intelligence and superior generalizability afforded by independent pre-trained transformer encoders for protein and ligand entities respectively permits the straightforward utilization of separated features that can subsequently be synthesized via downstream fully connected layers for affinity prediction.

For protein features, PLAPT incorporates ProtBERT, a BERT-based model, chosen for its exceptional ability in creating embeddings for use in downstream models, achieving excellent accuracy for secondary structure prediction, sub-cellular localization, and other tasks [17, 19]. Protein sequences, represented as token arrays (length 3200), as specified in Section 2.1, are processed through the ProtBERT model. The ensuing pooler output is a 1024-dimensional vector that contains essential features and structural information of the protein entity (see Figure 1 for visualization).

Correspondingly, PLAPT uses ChemBERTa for the extraction of ligand features. ChemBERTa, modeled on the RoBERTa transformer architecture specifically tailored for chemical data processing, produces molecular representations that preform relatively well for downstream molecular property prediction tasks, thus making it a good choice for use in PLAPT [22, 16]. The pooler output from ChemBERTa is a 768-dimensional vector that contains the molecular structure and relevant features of the ligand entity (see Figure 1 for visualization).

The resultant protein and molecule representations, of dimensions 1024 and 768 respectively, are concatenated into a composite protein-ligand feature vector of length 1792. This representation serves as the input for the succeeding prediction module, which utilizes a branching neural network, as visualized in Figure 1 and described in more detail in Section 3.

#### 2.2.2 Prediction Module

The Prediction Module in PLAPT is an BNN based regression model designed to compute protein-ligand binding affinities by processing the protein-ligand feature vector obtained from ProtBERT and ChemBERTa [19, 16]. The input to this module is the 1792-dimensional feature vector described in Section 2.2.1, which contains valuable information about both the protein and ligand. The feature vector is then split into bifurcated fully connected layers as part of our BNN architecture.

The prediction module processes a length 1792 feature vector in two streams: the first 1024 indices in a protein-specific layer and the remaining 768 in a molecule-specific layer, each followed by ReLU activation. The combined outputs undergo normalization and pass through three linear layers with ReLU activations and a dropout layer to reduce overfitting. The final output is a scalar value indicating the normalized negative log10 affinity. For more details, refer to Appendix 2.2.2, and for visualization, see Figure 3.

The rationale behind employing a Branching Neural Network architecture stems from the imperative to enhance the initial representations of proteins and ligands provided by ProtBERT and ChemBERTa. Although these representations are comprehensive, they are broad and require further refinement to optimally serve the purpose of affinity prediction. The BNN architecture facilitates this refinement by applying linear layers to further transform the protein and ligand features independently, thereby rendering them more suitable for downstream affinity prediction. This approach is predicated on the hypothesis that such transformations could improve the model’s predictive accuracy by tailoring the feature representations to be more relevant for affinity prediction tasks. This hypothesis aligns with existing literature that underscores the efficacy of feature transformation in enhancing machine learning tasks [23].

### 2.3 Datasets

The primary dataset for PLAPT was derived from a subset of the Binding Affinity Dataset, obtained from Glaser et al. via the Hugging Face platform [24]. This subset consisted of the first 100,000 data points, containing the amino acid sequence, canonical SMILES, and normalized negative logarithm (base 10) of affinity for each protein-ligand pair. The decision to use a reduced set was influenced by computational constraints.

Prior to model training, a dataset containing protein-ligand feature vectors was generated using the ProtBERT and ChemBERTa models, as described in Section 2.2.1. This process was executed on an NVIDIA RTX 2060 GPU, requiring approximately 12 hours to complete. Detailed compute specifications can be found in Appendix A.2. The resulting dataset was comprised of 100,000 instances of a 1792-dimensional feature vector paired with a binding affinity value, which would be subsequently used for training the prediction module. The complete feature dataset can be downloaded from our Google Drive linked in Section B.2.

Following the embedding generation, the data was split into training and validation sets using a seeded random split with a 90/10 ratio, resulting in 90,000 samples for the training set and 10,000 samples for the validation set. This process was done in Wolfram Language, the code for which being available in Section B.1.

For benchmarking purposes, we utilized the Test2016_290 dataset from Jin et al. [11], ensuring it had no overlap with the 100,000 data points designated for training and validation. We identified and corrected an erroneous double bond in the 263rd protein-ligand pair of the Test290 dataset, converting it to a single bond. Additionally, we used the CSAR-HiQ_36 dataset [25], which comprises 36 protein-ligand pairs and aligns with the study by Jin et al. [11]. Furthermore, to enhance our benchmarking resources, we created a new test dataset, herein called Benchmark1k2101. This dataset was generated by selecting 1,000 data points at random from indices 10001 to 20000 in the full Binding Affinity Dataset [24]. For reproducibility, we used the seed value 2101, consistent with the rest of our paper. All benchmark datasets are accessible on our GitHub repository, detailed in Appendix B.1.

### 2.4 Training

The prediction module underwent training using Wolfram Mathematica’s NetTrain function, known for its efficiency and ease-of-use [26]. We used a mean squared error loss function in conjunction with the Adam optimizer [27]. Training was performed on an Nvidia RTX 4060 ti 16GB, completing in approximately three minutes. For further details on the training hyperparameters and computational specifications, see Appendix A.4 and A.2 respectively.

### 2.5 Evaluation

To assess the efficacy of PLAPT in predicting protein-ligand binding affinities, we utilized three widely recognized metrics: the Pearson correlation coefficient (*R*), root mean square error (RMSE), and mean absolute error (MAE). The mathematical formulations of these metrics are delineated in Appendix A.3. The evaluation process involved a comparison between the experimental and predicted *pK*_*d*_ values, necessitating the denormalization of the model outputs prior to analysis.

We compared the performance of our model, PLAPT, against several other protein-ligand binding affinity predictors, including CAPLA, CAPLA-Pred [11], DeepDTA [28], DeepDTAF [10], Pafnucy [8], OnionNet [9], FAST [29], IMCP-SF [30], GLI [31], and the affinity prediction model made by Blanchard et al. [18], on benchmarks described in Section 2.3.

## 3 Results

In this section, we present and analyze the performance metrics of PLAPT. We begin by detailing the training, validation, and performance metrics on our test set, Benchmark1k2101. This is followed by a comparative analysis against state-of-the-art models on two independent benchmark datasets, Test-2016_290 and CSAR-HiQ_36, highlighting PLAPT’s performance and generalizability [11, 25].

### 3.1 Performance of PLAPT

Table 1 presents the performance metrics of the PLAPT model across different datasets: the training set, the validation set, and the Benchmark1k2101 dataset. The metrics evaluated include the Pearson correlation coefficient (R), the root mean square error (RMSE), and the mean absolute error (MAE), mentioned in Section 2.5. The model demonstrates strong performance on the Benchmark1k2101 dataset, achieving high R values (R = 0.883) alongside lower RMSE (0.851) and MAE (0.688) compared to the training set, indicating effective generalization to unseen data and a strong correlation between predicted and experimental binding affinity values. However, the model’s validation set performance, indicated by an R value of 0.683, hints at possible overfitting due to a noticeable decline in correlation as compared to the 0.886 R value for training. Nonetheless, when evaluating PLAPT’s validation accuracy against other methodologies like CAPLA [11], which recorded a validation RMSE (Root Mean Square Error) and MAE (Mean Absolute Error) of 1.338 and 1.034 respectively, PLAPT demonstrates significantly lower validation RMSE and MAE, meaning it is likely no-more overfit than the state-of-the-art.

**Table 1:** Performance Metrics of the PLAPT Model.

| Data Set | R | MSE | RMSE | MAE |
| --- | --- | --- | --- | --- |
| Train Data | 0.886 | 0.586 | 0.765 | 0.756 |
| Benchmark1k2101 | 0.883 | 0.851 | 0.922 | 0.688 |
| Validation Data | 0.683 | 1.466 | 1.211 | 0.949 |

In Figure 2, graphs a, b, and c, display the distributions of *pK*_*d*_ values for the training, validation, and Benchmark1k2101 datasets, respectively. Both the training and validation datasets exhibit a skew towards lower *pK*_*d*_ values. While the impact of this skew on the model’s performance is not explicitly analyzed, the model’s predictions mirror this distribution trend. Moreover, the broad dispersion of data points in panel B highlights the model’s reduced validation accuracy, as evidenced by a Pearson correlation coefficient (R) of 0.683.

**Figure 2:**
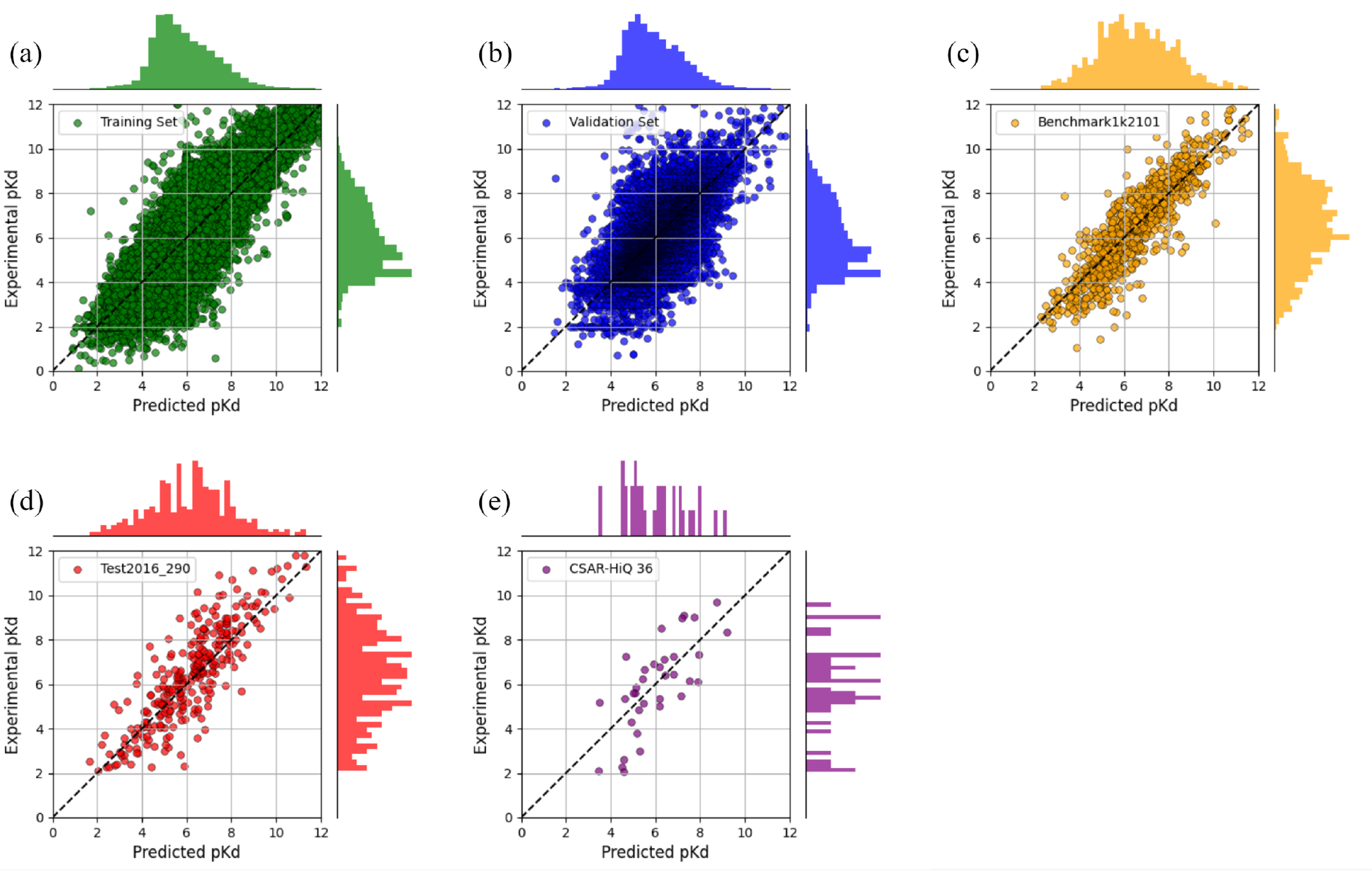
Evaluation of PLAPT on Multiple Datasets. Scatter plots depict the correlation between predicted and experimental *pK*_*d*_ values, while accompanying histograms display the distribution of both predicted and actual *pK*_*d*_ values. The datasets represented include the Training Set (a), Validation Set (b), Benchmark1k2101 Set (c), Test2016_290 (d), and CSAR-HiQ_36 (e)

**Figure 3:**
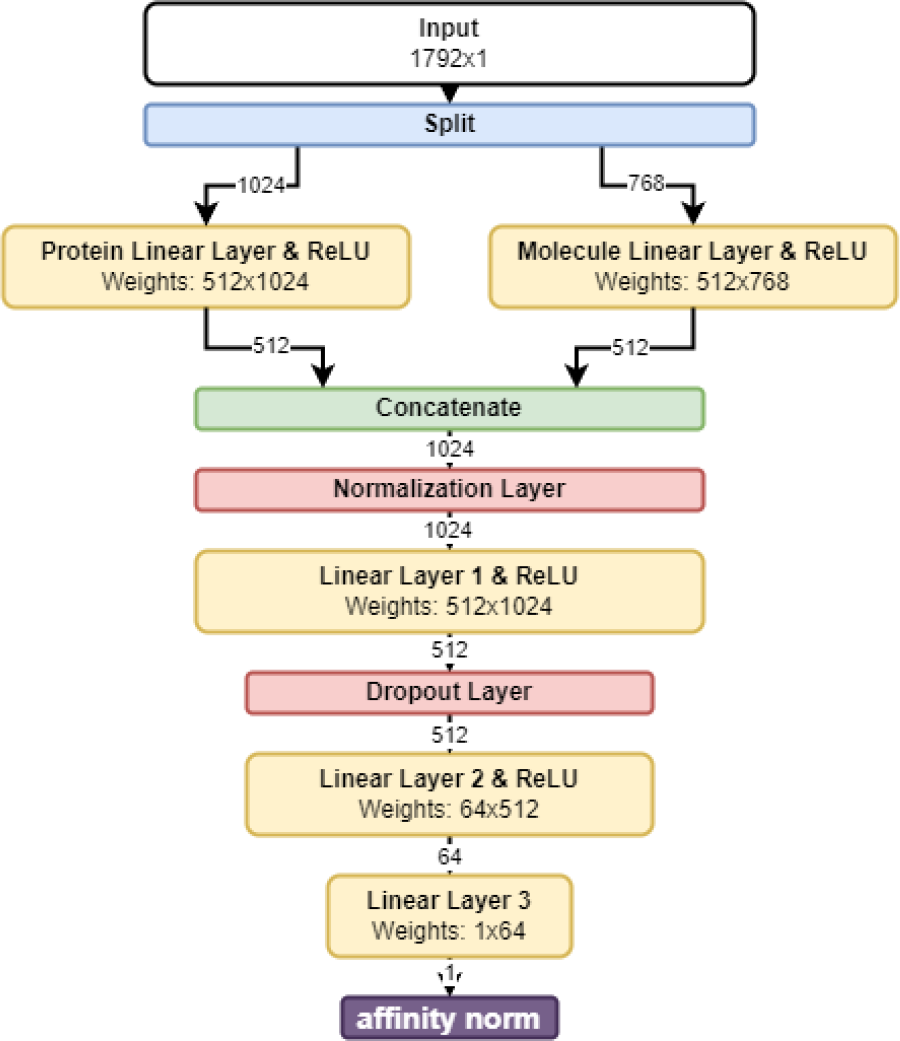
The prediction module takes a 1792×1 feature vector as its input, which is then partitioned into two streams: The first stream processes the first 1024 indices of the feature vector through a 512-node protein-specific linear layer, followed by a ReLU activation function. Concurrently, the second stream processes the latter 768 feature indices through a similar 512-node molecule-specific linear layer, also followed by a ReLU activation.

### 3.2 Comparative Analysis

To evaluate the performance of the PLAPT model, we conducted comparative analyses against other models using two distinct benchmark datasets: Test2016_290 [11] and CSAR-HiQ_36 [25], as described in Section 2.5. These comparisons aim to provide a comprehensive assessment of PLAPT’s efficacy in the context of current state-of-the-art methodologies.

#### Test2016_290 Benchmark

The Test2016_290 dataset, introduced in the study by Jin et al. [11], serves as a benchmark for evaluating the performance of PLAPT. As shown in Table 2, PLAPT demonstrates superior performance over competing models, including CAPLA, across several metrics: correlation coefficient (R), root mean square error (RMSE), and mean absolute error (MAE). However, it is important to note that the affinity prediction model by Blanchard et al. [18] outperformed PLAPT on all metrics for this benchmark. Subsequent analysis revealed that 177 out of the 290 data points in the Test2016_290 dataset [11] were also included in the training and validation dataset used by Blanchard et al., leading to their exclusion from this comparison due to potential overfitting concerns. PLAPT surpasses CAPLA [11] by 0.33%, CAPLA-Pred [11] by 7.86%, DeepDTA [28] by 17.12%, DeepDTAF [10] by 11.73%, Pafnucy [8] by 15.66%, OnionNet [9] by 6.42%, FAST [29] by 8.56%, IMCP-SF [30] by 17.63%, and GLI [31] by 7.57% for the RMSE metric. Additionally, PLAPT surpasses CAPLA [11] by 6.21%, CAPLA-Pred [11] by 10.65%, DeepDTA [28] by 21.08%, DeepDTAF [10] by 15.56%, Pafnucy [8] by 19.75%, FAST [29] by 11.09%, IMCP-SF [30] by 21.56%, and GLI [31] by 11.70% for the MAE metric.

**Table 2:** Comparison of PLAPT with Various Other Methods [11, 28, 10, 8, 9, 29, 30, 31] on the Test2016_290 Benchmark [11].

| Method | R | RMSE | MAE |
| --- | --- | --- | --- |
| PLAPT | **0.845** | **1.196** | **0.906** |
| CAPLA | 0.843 | 1.200 | 0.966 |
| CAPLA-Pred | 0.825 | 1.298 | 1.014 |
| DeepDTA | 0.749 | 1.443 | 1.148 |
| DeepDTAF | 0.789 | 1.355 | 1.073 |
| Pafnucy | 0.775 | 1.418 | 1.129 |
| OnionNet | 0.816 | 1.278 | – |
| FAST | 0.810 | 1.308 | 1.019 |
| IMCP-SF | 0.791 | 1.452 | 1.155 |
| GLI | – | 1.294 | 1.026 |

#### CSAR-HiQ_36 Benchmark

In an analysis of model performance on CSAR-HiQ_36 [25], PLAPT demonstrated superior performance across multiple metrics, including Pearson’s correlation coefficient (R), root mean square error (RMSE), and mean absolute error (MAE), as compared to other models such as CAPLA, DeepDTAF, Pafnucy, IGN, and IMCP-SF [11, 10, 8, 32, 30]. This is evidenced by the data presented in Table 3. However, the affinity_pred model by Blanchard et al. achieved a higher Pearson’s correlation coefficient (R = 0.774) compared to PLAPT (R = 0.731), indicating a slightly better performance in capturing the linear relationship between predicted and actual binding affinities. Despite this, PLAPT’s performance is more accurate than affinity_pred [18] on this benchmark, with a 9.10% improvement in RMSE and a 1.62% improvement in the MAE metric. Aside from affinity_pred, PLAPT surpasses CAPLA [11] by 7.22%, DeepDTAF [10] by 51.21%, Pafuncy [8] by 18.64%, IGN [32] by 24.85%, and IMCP-SF [30] by 13.53% for the RMSE metric. For the MAE metric, PLAPT surpasses CAPLA [11] by 0.26%, DeepDTAF [10] by 50.09%, Pafnucy [8] by 10.38%, IGN [32] by 19.15%, and IMCP-SF [30] by 3.98%.

**Table 3:** Comparison of PLAPT with Various Methods on the CSAR-HiQ_36 Dataset.

| Method | R | RMSE | MAE |
| --- | --- | --- | --- |
| PLAPT | 0.731 | **1.349** | **1.157** |
| affinity_pred | **0.774** | 1.484 | 1.176 |
| CAPLA | 0.704 | 1.454 | 1.160 |
| DeepDTAF | 0.543 | 2.765 | 2.318 |
| Pafnucy | 0.566 | 1.658 | 1.291 |
| IGN | 0.528 | 1.795 | 1.431 |
| IMCP-SF | 0.631 | 1.560 | 1.205 |

**Table 4:** Specifications of the computers used for the encoding and the training processes.

| Specification | Encoding | Training |
| --- | --- | --- |
| CPU | Intel i7-9700K | Intel i7-13700KF |
| GPU | NVIDIA RTX 2060 | NVIDIA RTX 4060-ti |
| RAM | 32 GB | 32 GB |
| VRAM | 6 GB | 16 GB |
| Operating System | Windows 11 | Windows 11 |

**Table 5:** List of training hyperparameters.

| Hyperparameter | Value |
| --- | --- |
| Loss Function | MSE |
| Optimizer | Adams |
| Learning Rate | 0.001 |
| Batch Size | 256 |
| Max Training Rounds | 60 |
| Random Seed | 2101 |

## 4 Conclusion

In this study, we introduced the Protein Ligand Binding Affinity Prediction Using Pre-trained Transformers (PLAPT) model, a novel machine learning approach aimed at accurately predicting protein-ligand binding affinities with significantly reduced computational resources. Our findings demonstrate that PLAPT achieves state-of-the-art accuracy, outperforming existing models across various benchmarks. This success is attributed to the innovative use of pre-trained transformers, specifically ProtBERT and ChemBERTa, integrated through a branching neural network architecture to efficiently predict binding affinities from one-dimensional sequence data.

The significance of PLAPT extends beyond its predictive performance. Accurate prediction of protein-ligand binding affinities is crucial in drug discovery and design, as it enables the rapid identification of promising drug candidates, thereby conserving valuable time and resources. By leveraging pre-trained models, PLAPT minimizes the need for extensive computational power for training, making the development of new predictive capabilities more accessible to researchers and institutions with limited resources.

Despite its strengths, PLAPT is not without limitations. The use of a reduced dataset, driven by computational constraints, and the reliance on a simplified model architecture suggest areas for potential improvement. Future research could explore the incorporation of binding site data to enhance the model’s predictive capabilities or investigate alternative architectures such as an attention-based prediction module to further improve accuracy.

Looking ahead, there is significant potential for expanding the scope of PLAPT. Future work could involve the exploration of more extensive datasets, the integration of binding site data, and the validation of the model across a broader range of datasets. These efforts would not only test the robustness and generalizability of PLAPT but also contribute to a deeper understanding of molecular interactions, paving the way for the next generation of drug discovery models.

In conclusion, PLAPT represents a significant advancement in the prediction of protein-ligand binding affinities. Its innovative use of pre-trained transformers, combined with a focus on efficiency and accessibility, positions PLAPT as a valuable tool in the field of drug discovery and design. As we continue to explore the potential of machine learning and deep learning in biotechnology, models like PLAPT underscore the transformative impact of computational approaches, promising to accelerate the pace of discovery and innovation in the search for new therapeutic agents.

## Acknowledgements

This project was conducted in part using Wolfram Language as a component of the Wolfram Emerging Leaders Program. We extend our gratitude to the Wolfram team for their support and for providing the resources that helped facilitate our research.

## A. Appendix

### A.1 Prediction Module Details

Outputs from both streams are concatenated into a single vector of 1024 elements. This combined vector is passed through a batch normalization layer with a momentum of 0.9 and epsilon of 0.001. The vector is then fed through the 512-node Linear Layer 1 and ReLU activation. A dropout layer with a probability of 20% is then applied to mitigate overfitting.

Following the dropout layer, the prediction module continues to reduce the feature space with the 64-node Linear Layer 2 with ReLU activation, before reaching the single node Linear Layer 3. This layer outputs the predicted scalar value representing the normalized negative log10 affinity value, which is then un-normalized.

### A.2 Compute

Two computers were used to create the PLAPT model. One was used to obtain the feature dataset, as described in Section 2.3. Another was used for model training, as described in Section 2.4.

### A.3 Definition of Performance Metrics

In this section, we define the performance metrics used to evaluate PLAPT.

#### Pearson’s Correlation Coefficient

The Pearson’s correlation coefficient (*r*) measures the linear correlation between the observed and predicted values. It is defined as:

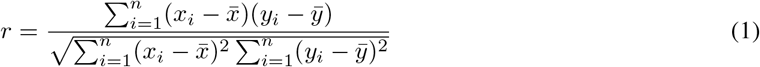

where *x*_*i*_ and *y*_*i*_ are the observed and predicted values, respectively, 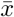 and 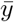 are their respective means, and *n* is the number of data points.

#### Root Mean Square Error (RMSE)

The RMSE is a measure of the differences between values predicted by a model and the values actually observed. It is defined as:

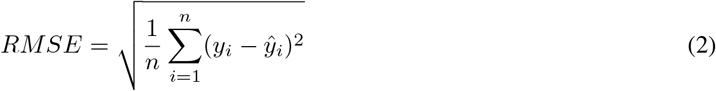

Here, *y*_*i*_ represents the observed values, *ŷ*_*i*_ represents the predicted values, and *n* is the number of observations.

#### Mean Absolute Error (MAE)

The MAE measures the average magnitude of the errors in a set of predictions, without considering their direction. It is defined as:

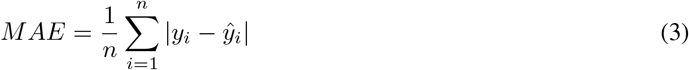

In this equation, *y*_*i*_ and *ŷ*_*i*_ again represent the observed and predicted values, respectively, and *n* is the number of observations.

### A.4 Training Details

Training was done in Wolfram Language (Mathematica), specifically the NetTrain function [26]. Training code can be found on PLAPT’s GitHub repository in Appendix B.1.

## B. Supplementary Materials

In this section, we provide links to the code and datasets used in our study.

### B.1 Code Repository

The source code for training, inference, and benchmark dataset files, is available on PLAPT’s GitHub repository. It can be accessed at the following URL:

https://github.com/trrt-good/WELP-PLAPT/tree/main

### B.2 Feature Dataset

The feature dataset used for training, supplementary model files, and tokenizer information, is hosted on Google Drive. It can be accessed using the link below:

https://drive.google.com/drive/folders/1e-ujgHx5bW0JKxSZY5u34As77o4-IIFs?usp=sharing

## Notes

https://github.com/trrt-good/WELP-PLAPT/tree/main

### Competing Interest Statement

This project was conducted in part using Wolfram Language, which was granted as a component of the Wolfram Emerging Leaders Program.

### Summary of Updates

citation fix typo fix orcid id

